# Label-free classification of cell death pathways via holotomography-based deep learning framework

**DOI:** 10.1101/2025.06.06.658404

**Authors:** Minwook Kim, Wei Sun Park, Geon Kim, Sanggeun Oh, Jaephil Do, Juyeon Park, YongKeun Park

## Abstract

Accurate classification of cell death pathways is critical in understanding disease mechanisms and evaluating therapeutic responses, as dysregulated cell death underlies a wide range of pathological conditions including cancer and therapy resistance. Conventional imaging methods such as fluorescence and bright-field microscopy, or 2D phase imaging, often suffer from phototoxicity, labeling artifacts, or limited morphological contrast. Here, we present a real-time, label-free platform for classifying cell death phenotypes—apoptosis, necroptosis, and necrosis—by combining three-dimensional holotomography with deep learning. Our convolutional neural network, trained on refractive index (RI)-based features from HeLa cells, achieved high classification accuracy (97.2 ± 2.8%) under varying cell densities. Notably, the model identified early RI changes during necroptosis several hours prior to fluorescence-based markers. These findings demonstrate the potential of holotomography-based AI for high-resolution, label-free cell death profiling.

## 1. Introduction

Cell death is a fundamental biological process essential for maintaining tissue homeostasis and regulating organismal development^1^. Dysregulation of cell death underlies a wide range of pathological conditions, including cancer, neurodegeneration, and autoimmune disorders, where inappropriate activation or evasion of death pathways contributes to tumor progression, immune escape, and tissue damage. Accurate and timely identification of distinct cell death modalities— particularly apoptosis, necroptosis, and necrosis—is therefore critical for advancing both mechanistic understanding and therapeutic decision-making in biomedical research and clinical applications^2-4^.

Fluorescence microscopy has long been the gold standard for cell death classification due to its high chemical specificity through targeted staining of intracellular markers (Fig. 1a). In recent years, machine learning (ML) and deep learning (DL) methods have been increasingly employed to automate such analyses based on fluorescence-labeled morphological and intensity features^5-7^. However, these techniques rely heavily on staining, fixation, and other invasive procedures that limit temporal resolution and preclude label-free or long-term live-cell imaging. Furthermore, the use of phototoxic dyes and photobleaching limits their applicability for dynamic phenotyping.

**Figure 1:**
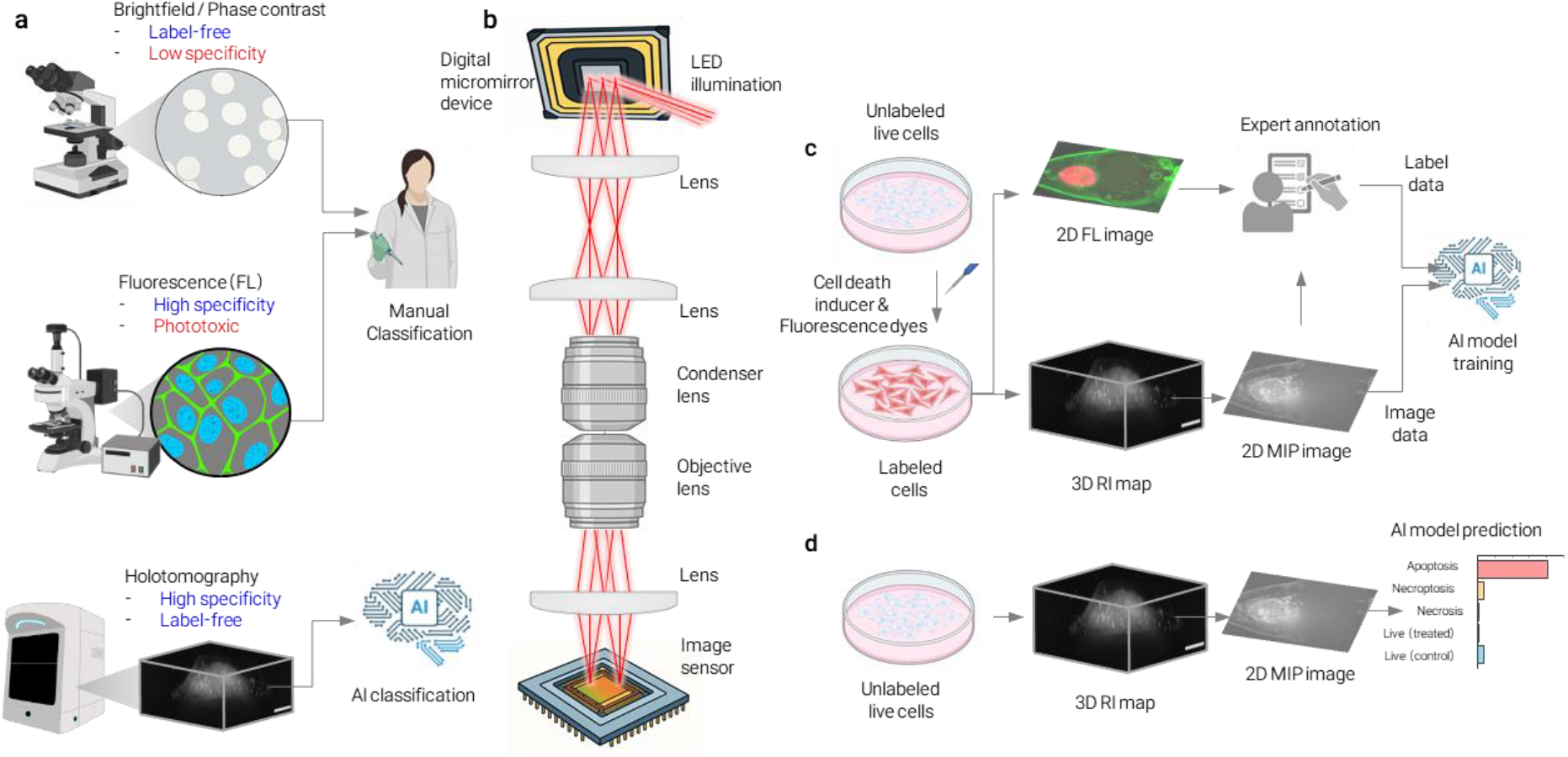
AI-powered, label-free classification of cell death pathways using holotomography. a, Comparison of conventional live-cell imaging modalities. Bright-field microscopy offers label-free imaging but suffers from low specificity. Fluorescence microscopy provides high chemical specificity but introduces phototoxicity and photobleaching, limiting its suitability for long-term live-cell monitoring. Holotomography addresses these limitations by enabling high-resolution, label-free, and non-invasive 3D imaging. b, Schematic of the holotomography system. Incoherent light from an LED source is patterned via a digital micromirror device (DMD) and projected onto the sample at multiple angles. The resulting diffraction patterns are recorded across various axial positions and computationally reconstructed into 3D refractive index (RI) tomograms using a tomographic inversion algorithm. c, Overview of the AI training pipeline. Cells are chemically treated to induce distinct death pathways and co-registered fluorescence images are used to generate expert-verified ground-truth annotations. Corresponding 3D RI tomograms are converted to 2D maximum intensity projections (MIPs) for use in deep learning model training. d, Inference phase. The trained model is applied to untreated or unseen test samples to perform label-free classification of cell death phenotypes in real time.

To overcome these limitations, various label-free optical imaging modalities have been explored. While phase contrast microscopy has been used for visualizing live cells without staining, its qualitative nature limits its utility in automated live/dead assays. More recently, quantitative phase imaging (QPI) has emerged as a promising alternative, offering precise and reproducible measurements of optical phase delay that reflect cellular morphology and biophysical status^8-11^. The quantitative nature of QPI provides a significant advantage when training deep learning models, as it enables the extraction of consistent and information-rich features^12-15^. However, despite this progress, conventional 2D QPI techniques are fundamentally limited by shallow depth resolution and the inability to capture subcellular structures critical for distinguishing morphologically similar cell death pathways^16-18^.

In particular, existing label-free approaches are often limited by one or more of the following: (i) reliance on 2D projections lacking volumetric context^16,17^, (ii) restricted classification to binary or coarse states (e.g., live vs. dead)^17,18^, and (iii) limited generalizability across different cell types, stimuli, or death mechanisms^11,16,18^. Necroptosis—a regulated form of necrosis that shares features with both apoptosis and classical necrosis—exemplifies this challenge due to its overlapping morphology and temporal dynamics.

To address these challenges, we present a deep learning framework that integrates holotomography— a three-dimensional (3D), label-free quantitative phase imaging technique—with convolutional neural networks (CNNs) to classify cell death phenotypes. Holotomography (HT) reconstructs volumetric refractive index (RI) maps from multiple 2D holographic measurements with various illumination or sample conditions, enabling non-invasive, high-resolution imaging of subtle biophysical changes in live cells (Fig. 1b)^19-24^. These RI-based features offer rich, label-free morphological contrast that can be leveraged for data-driven classification^25-27^.

From a machine learning perspective, cell death classification using 3D RI maps poses a unique yet underexplored challenge: How can models reliably detect subtle, heterogeneous, and temporally evolving morphological changes? And how well can these models generalize across domains—such as different cell types, death inducers, or treatment conditions—that exhibit variation in morphological expression?

In this work, we present a CNN-based classification model trained on MIP mages derived from 3D RI tomograms to identify five cell states: apoptosis, necroptosis, necrosis, untreated live, and drug-treated live cells. Using a curated dataset of co-registered RI and fluorescence images from HeLa cells, we fine-tune a ResNet-101 architecture to perform patch-wise inference without requiring explicit single-cell segmentation. Our model achieves high accuracy (97.2 ± 2.8%) and performs robustly in densely packed, wide-field imaging conditions. We further evaluate the model’s generalizability using A549 lung cancer cells, where performance degradation due to domain shift highlights critical limitations of morphology-only classification. Overall, this work demonstrates the potential of holotomography-based AI for high-resolution, real-time, label-free phenotyping of cell death, with broad implications for drug screening, cytotoxicity testing, and dynamic cell fate analysis.

## 2. Results

### 2.1 Morphological signatures of cell death pathways revealed by 3D holotomography

To evaluate the potential of RI-based label-free classification of cell death phenotypes, we acquired high-resolution 3D RI tomograms of HeLa cells subjected to four distinct experimental conditions: untreated control, apoptosis (induced by FasL or TNFα), necroptosis (induced by TNFα with z-VAD), and necrosis (induced by NaOH treatment) (Figs. 2a–d). HT reconstructions revealed distinct volumetric morphologies characteristic of each condition, establishing a structural basis for downstream classification.

**Figure 2:**
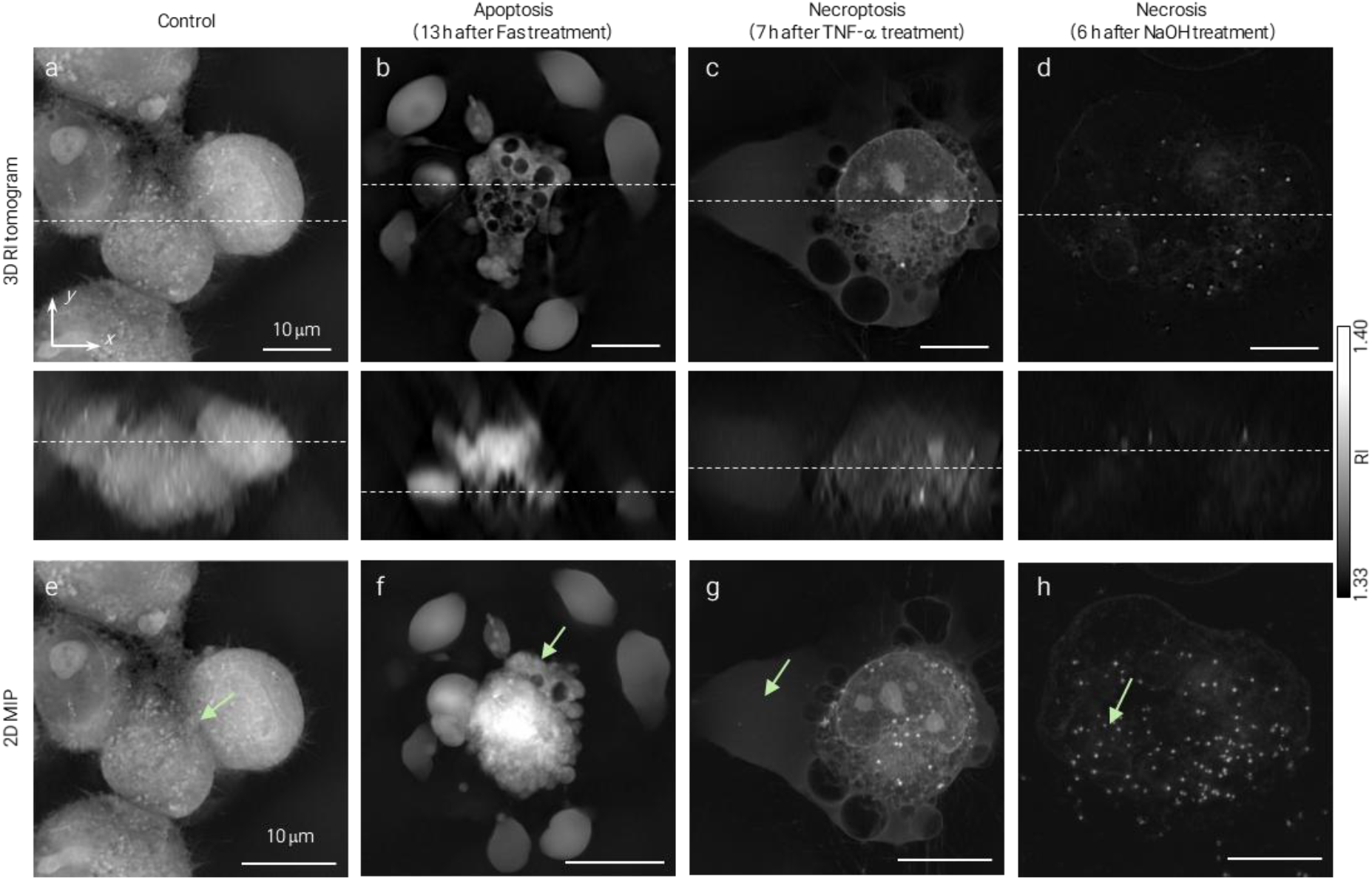
Label-free morphological features of distinct cell death pathways captured by 3D holotomography. a–d, Representative refractive index (RI) tomograms of HeLa cells under four different conditions: a, Live (untreated control); b, Apoptosis induced by Fas ligand (FasL); c, Necroptosis induced by TNF-α in combination with the pan-caspase inhibitor z-VAD; d, Necrosis induced by sodium hydroxide (NaOH). Each panel presents orthogonal RI slices in the x–y plane (top row) and x–z plane (bottom row), highlighting condition-specific 3D morphological features. e–h, Corresponding 2D maximum intensity projection (MIP) images for each condition. Green arrows indicate representative hallmarks: e, Mitotic control cells with intact membrane and nuclear structure; f, Apoptotic cells showing membrane blebbing and cytoplasmic condensation; g, Necroptotic cells with expanded cell volume and preserved nuclear morphology; h, Necrotic cells displaying diffuse RI loss and granular texture, consistent with cytoplasmic leakage.

Control cells exhibited intact plasma membranes and well-defined nuclear boundaries, often undergoing mitotic division (Fig. 2e), consistent with healthy and proliferative morphology. Apoptotic cells showed classical features of programmed cell death, including shrinkage, membrane blebbing, and the formation of apoptotic bodies (Fig. 2f), in agreement with known caspase-mediated mechanisms^28-30^.Necroptotic cells were characterized by increased cell volume and preserved nuclear structure (Fig. 2g), clearly distinguishing them from necrotic cells. Despite signs of cytoplasmic remodeling, RI values in necroptotic cells remained relatively high, indicating membrane integrity was still maintained. In contrast, necrotic cells displayed a diffuse and markedly reduced RI distribution, reflecting extensive intracellular leakage and membrane breakdown (Fig. 2h)^31^. A homogeneous drop in RI and the presence of granular cytoplasmic structures emerged as consistent necrotic signatures in HT images.

These results demonstrate that HT provides sufficient biophysical contrast to differentiate morphologically overlapping but mechanistically distinct forms of cell death. In particular, the ability to distinguish necroptosis from apoptosis based on volumetric structure and RI preservation supports the feasibility of morphology-driven, label-free classification—and serves as the foundation for the deep learning model developed in this study.

### 2.2 Deep learning–based classification of cell death using RI-derived projections

To develop a model capable of classifying cell death phenotypes from label-free HT images, we constructed a dataset of 3D RI tomograms acquired 16 hours post-treatment—a time point when cell death–associated morphological features were stabilized. Each 3D RI volume was converted into a 2D MIP along the z-axis to reduce computational complexity while preserving salient morphological information. The MIP images were subdivided into fixed-size patches (320 × 320 pixels; 51.2 μm per side), and non-cell regions were excluded via RI thresholding (see Methods).

The resulting dataset comprised five classes: apoptosis (n = 91), necroptosis (n = 136), necrosis (n = 43), live-control (n = 104), and live-treated (n = 159). An independent test set was constructed using samples from separate culture batches (n = 21, 34, 11, 26, and 55 for each class, respectively). We employed a transfer learning strategy using a ResNet-101 model pretrained on ImageNet (Fig. 3a), fine-tuning the final convolutional and fully connected layers to accommodate five-class classification. This enabled efficient adaptation to cell morphology–specific features.

**Figure 3:**
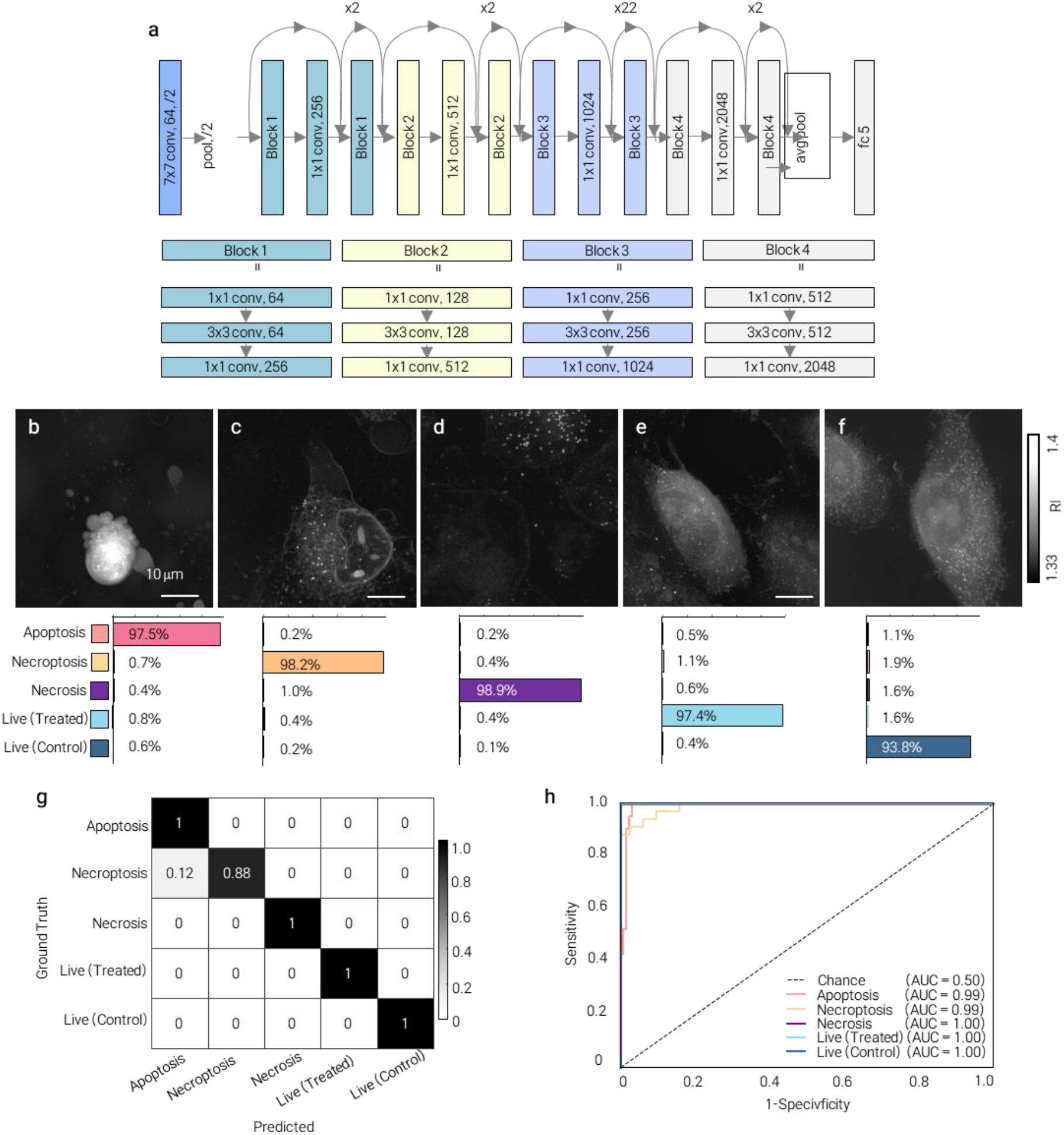
Deep learning–based classification of cell death phenotypes using holotomography-derived image patches. a, Architecture of the ResNet-101 convolutional neural network employed in this study. The model was fine-tuned from an ImageNet-pretrained backbone by modifying and retraining the final convolutional and fully connected layers to classify five distinct cell states. b–f, Representative 2D MIP patches (320 × 320 pixels; 51.2 μm per side) corresponding to each class: b, apoptosis; c, necroptosis; d, necrosis; e, live-treated; f, live-control. Softmax probability bars below each panel illustrate the model’s predicted confidence distribution over all classes. Despite morphological heterogeneity, the model produces high-confidence predictions across phenotypically diverse cell states. g, Confusion matrix summarizing classification performance across five classes— apoptosis, necroptosis, necrosis, live-control, and live-treated—based on patch-wise predictions. The model achieved an overall accuracy of 97.2 ± 2.8%, with the primary confusion occurring between necroptosis and apoptosis (12% misclassification). h, Receiver operating characteristic (ROC) curves illustrating class-wise model performance, with sensitivity (true-positive rate) plotted against 1– specificity (false-positive rate).

Representative MIP patches from each class, along with predicted softmax probability distributions, are shown in Figs. 3b–f. Despite variability in cell morphology and confluency, the model consistently assigned high-confidence predictions, demonstrating that patch-level RI features are sufficient for phenotype discrimination—even without segmentation or temporal context.

The trained model achieved an overall accuracy of 97.2 ± 2.8% on the test set (Fig. 3g). Most classes were predicted with high confidence, although 12% of necroptotic patches were misclassified as apoptotic—likely reflecting biological and morphological overlap between the two pathways, especially during transitional states^32^.

To further evaluate the model’s class-wise discriminative performance, we computed receiver operating characteristic (ROC) curves for each class based on patch-level predictions (Fig. 3g). All five cell states exhibited high area under the ROC curve (AUC) values, indicating excellent separability (Fig. 3h). Apoptosis, necrosis, and live-treated classes showed near-perfect AUCs approaching 1.0, while necroptosis and live-control exhibited slightly lower values due to partial overlap with adjacent classes—consistent with trends observed in the confusion matrix.

These findings confirm that the model’s probabilistic outputs are well-calibrated and that refractive index–derived features support high-sensitivity, high-specificity classification across morphologically diverse cell death phenotypes. Together, these results validate the feasibility of supervised learning for label-free, morphology-based cell death classification. The patch-wise design further enables scalable inference in densely populated and heterogeneous imaging contexts, laying the groundwork for wide-field and time-resolved applications.

### 2.3 Wide-field inference and time-resolved monitoring of cell death progression

To assess the scalability and practical utility of the model, we applied it to a wide-field stitched MIP image (3,796 × 3,796 pixels; 588 μm per side) comprising approximately 150 HeLa cells (Fig. 4a). This dataset was acquired from an independent experimental batch and excluded from both training and testing, providing an unbiased evaluation of model performance under realistic, high-confluency conditions.

**Figure 4:**
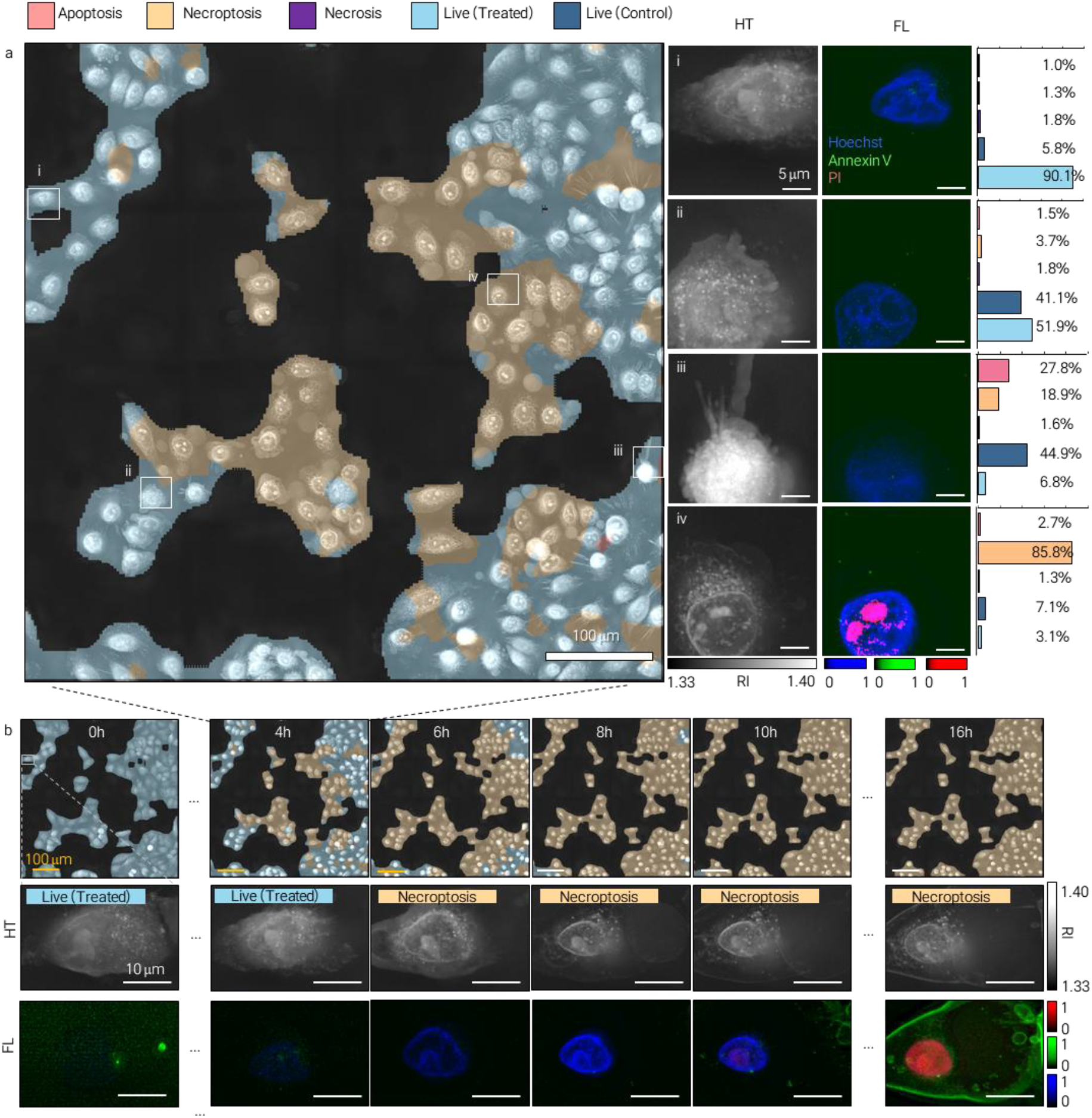
Time-resolved, label-free classification of cell death phenotypes in wide-field holotomography. a, Large-area 2D MIP image (588 μm × 588 μm) of a HeLa cell population (∼150 cells), acquired from a sample excluded from model training and testing. Each patch (160 × 160 pixels; 25.6 μm per side) was classified by the AI model into one of five categories—apoptosis, necroptosis, necrosis, live-control, and live-treated—based on refractive index (RI) features, and visualized using color-coded overlays. i–iv, Representative regions from panel a, showing RI images (left) and co-registered fluorescence channels (Hoechst: blue, Annexin V: green, PI: red). Bars indicate the model-predicted class probabilities. b, Time-lapse prediction results over a 16-hour imaging period. Patch-wise classification maps (top row) reveal a progressive population-level transition from live-treated (light blue) to necroptotic (orange) states. A representative single cell (bottom row) shows RI-based morphological changes preceding Annexin V and PI signal uptake in fluorescence channels (bottom row), demonstrating that the model predicts necroptosis approximately 4–6 hours earlier than biochemical markers. Scale bars: 100 μm in a, b (top); 10 μm in b (middle and bottom); 5 μm in i–iv.

We employed a patch-overlapping inference strategy using a sliding window (160 × 160 pixels; 25.6 μm per side), followed by majority voting across overlapping regions to enhance spatial consistency and mitigate boundary artifacts. The model accurately predicted cell death phenotypes across the field, maintaining robust performance even in densely populated regions with confluency up to 70%.

To assess prediction accuracy, we extracted representative patches from multiple regions and compared model outputs with co-registered fluorescence markers (Hoechst, Annexin V, and PI; Fig. 4i–iv). The strong spatial agreement between predicted labels and fluorescence-based ground truth supports the reliability of our model in dense, unsegmented settings.

We further investigated the model’s capacity to monitor dynamic cell state transitions using time-lapse imaging. At the population level, we observed a temporal shift from predominantly live-treated cells to necroptotic phenotypes over a 16-hour period (Fig. 4b, top). This trend was also evident at the single-cell level: one representative cell exhibited progressive morphological changes characteristic of necroptosis—namely, cell swelling, nuclear preservation, and eventual RI reduction (Fig. 4b, bottom).

Remarkably, the model predicted necroptotic transition approximately 4–6 hours earlier than detection by conventional fluorescence markers such as Annexin V and PI (Fig. 4b, bottom row). This observation highlights a key advantage of holotomography-based AI: the ability to identify early, label-free morphological cues of regulated cell death prior to biochemical signal emergence^33^.

Collectively, these results demonstrate that our model enables real-time, segmentation-free, and spatially resolved tracking of cell death dynamics at both population and single-cell levels. This capability is particularly valuable for high-throughput drug screening and cytotoxicity profiling, where label-free, time-resolved, and scalable classification is essential.

### 2.4 Generalization to A549 cells reveals domain-shift limitations

To assess the generalizability of our trained model, we applied the model trained with HeLa cells to A549 cells—a human lung adenocarcinoma line that is morphologically distinct from HeLa cells. As in the HeLa dataset, we induced three cell death modalities (apoptosis, necroptosis, and necrosis), along with live-control and drug-treated conditions. However, patches from cells treated with FasL + CHX + z-VAD—originally designed to induce necroptosis—were excluded from evaluation due to inconsistent and ambiguous morphologies (Fig. 5a).

**Figure 5:**
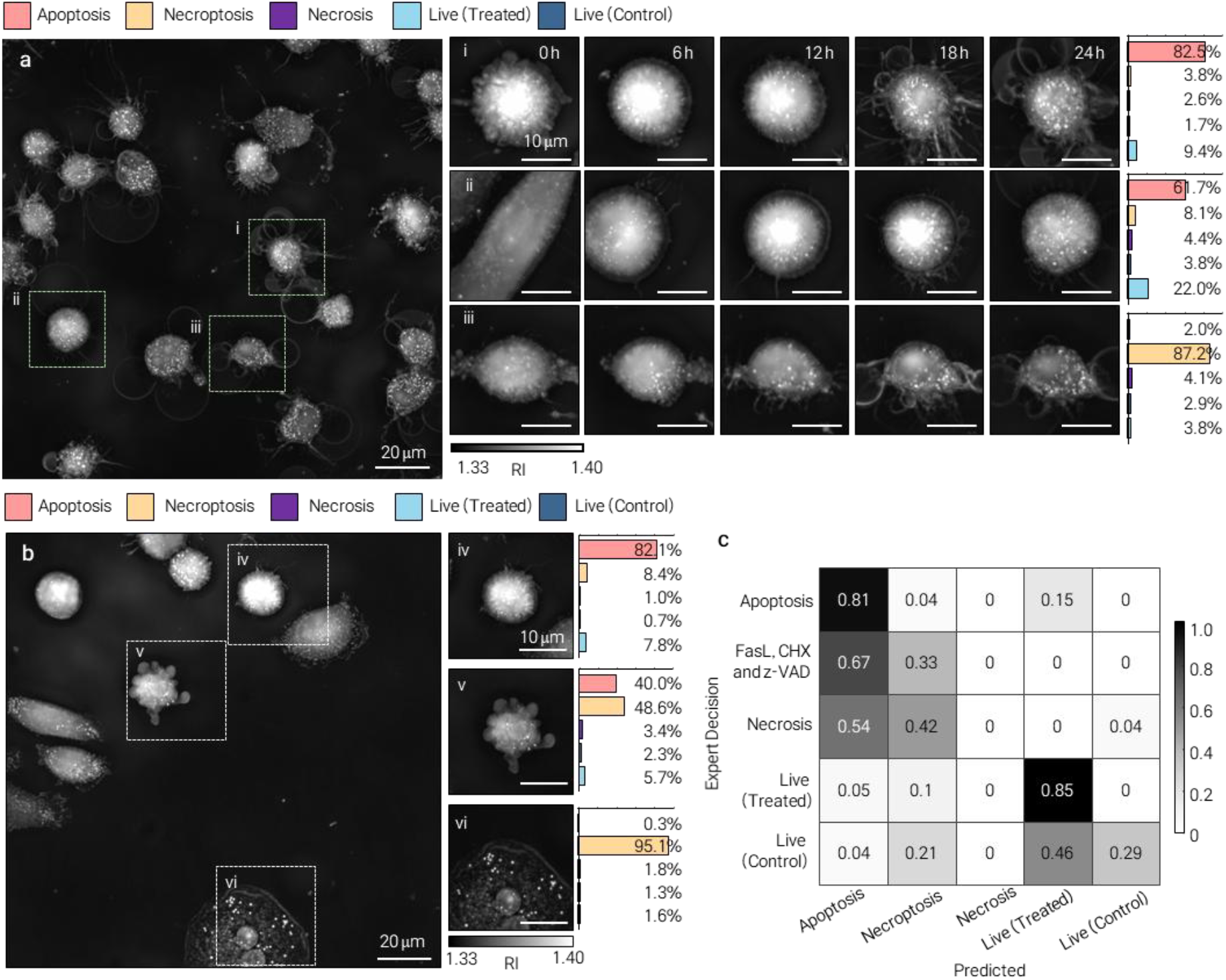
Heterogeneous cell death phenotypes in A549 cells and generalizability of the classification model. a, MIP image of A549 cells at 24 hours after treatment with Fas ligand (FasL), cycloheximide (CHX), and z-VAD. Green boxes indicate regions shown in panels i–iii. Each inset shows a high-resolution RI image of an individual cell, accompanied by a time-lapse series depicting morphological changes over 24 hours. Despite the treatment being designed to induce necroptosis, both apoptotic-like (i, ii) and necroptotic-like (iii) features are observed. Notably, these A549 cells exhibit progressive shrinkage and membrane rupture without nuclear preservation, contrasting with canonical necroptosis in HeLa cells and suggesting the presence of secondary necrosis. b, Inference results on a wide-field MIP image of A549 cells (unseen during training). The image was divided into 320 × 320 pixel patches and classified using the pretrained model. Boxes iv–vi show representative RI patches and predicted class probabilities for apoptosis (iv), necroptosis (v), and necrosis (vi) based solely on label-free input. c, Confusion matrix comparing AI predictions against expert annotations for A549 cells, after excluding morphologically ambiguous FasL + CHX + z-VAD-treated samples. While apoptosis and live-treated cells were classified with high accuracy, predictions for necrosis and live-control showed substantial degradation—likely due to morphological domain shift between A549 and HeLa cells. Scale bars: 20 μm in a, b; 10 μm in i–vi. RI values range from 1.33 to 1.40.

Unlike HeLa cells, A549 cells under necroptosis-inducing conditions frequently lacked defining features such as preserved nuclear structure and uniform cytoplasmic expansion. Time-lapse imaging revealed a sequential progression from apoptotic-like features (e.g., membrane blebbing, nuclear condensation) to loss of membrane integrity and diffuse RI dispersion, consistent with secondary necrosis (Fig. 5b). These hybrid transitions challenged definitive phenotype assignment, and in the absence of biochemical validation (e.g., RIPK3 or MLKL activation), classification as necroptosis could not be confidently established.

After excluding these ambiguous samples, model performance was evaluated on the remaining A549 classes. The overall classification accuracy dropped to 58%, highlighting a significant generalization gap (Fig. 5c). While apoptotic (81%) and live-treated (85%) cells were correctly classified, necrotic cells induced by hydrogen peroxide (H_2_O_2_) were completely misclassified (0%), and live-control cells were frequently confused with treated-live cells (46% misclassified).

This failure likely stems from two compounding factors. First, the model was trained exclusively on NaOH-induced necrosis in HeLa cells, characterized by rapid and homogeneous RI loss. In contrast, H_2_O_2_-treated A549 cells exhibited more subtle and spatially heterogeneous RI reductions, along with granular cytoplasmic textures (Fig. 6). Second, the morphological differences between live-control and treated A549 cells were subtler than in HeLa cells, and this intra-class variation was not sufficiently represented in the training set.

**Figure 6:**
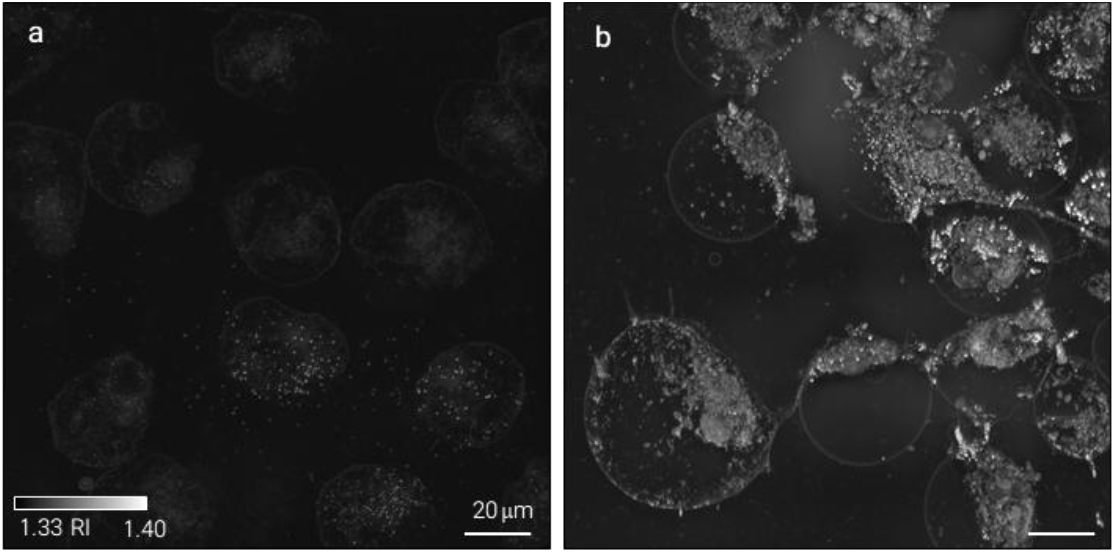
Label-free comparison of necrotic morphologies in HeLa and A549 cells using holotomography. a, HeLa cells treated with sodium hydroxide (NaOH) exhibit features of alkali-induced necrosis, including a homogeneously reduced refractive index (RI) across the cytoplasm, consistent with extensive membrane rupture and cytoplasmic leakage. b, In contrast, A549 cells subjected to hydrogen peroxide (H_2_O_2_) display oxidative necrosis, characterized by spatially heterogeneous RI patterns and prominent intracellular granularity, rather than diffuse RI loss. Both images were rendered using the same linear RI scale (1.33–1.40) to enable direct visual comparison. Scale bars: 20 μm.

These findings illustrate the challenge of domain shift in morphology-based classification. Despite strong performance on the source domain, even modest differences in cell type, death mechanism, or treatment condition can significantly impair generalization. Addressing this limitation will require more diverse and representative training datasets, incorporation of biochemical markers, or adoption of domain adaptation strategies to improve robustness in cross-context applications.

## 3. Discussion

In this study, we developed an integrated platform that combines 3D HT with DL to enable real-time, label-free classification of cell death pathways in live cells. Unlike conventional fluorescence-based assays—which require staining, fixation, and are subject to phototoxicity and bleaching—our approach allows continuous, non-invasive monitoring of dynamic morphological changes associated with apoptosis, necroptosis, and necrosis. By leveraging RI–based volumetric imaging and AI-driven pattern recognition, this method provides a powerful alternative to fluorescence microscopy for investigating regulated cell death in its native, label-free context.

Our findings highlight two key advantages of HT for cell death analysis. First, necrotic cells exhibit a pronounced and spatially uniform decrease in RI, resulting from cytoplasmic leakage and loss of membrane integrity (Fig. 2h). This RI-based contrast provides a distinct optical signature that facilitates label-free detection of necrosis—surpassing the sensitivity and interpretability of conventional fluorescence, bright-field, or phase contrast imaging^31^. Second, our model was able to identify early RI changes characteristic of necroptosis well before the appearance of conventional fluorescence markers such as Annexin V and PI (Fig. 4b). This temporal advantage is consistent with prior observations from digital holographic microscopy, where phase-based signals preceded biochemical signs of membrane rupture by several hours^33^. These results suggest that HT may uncover previously inaccessible early morphological cues associated with programmed cell death, offering new opportunities for time-resolved phenotyping and mechanism-driven screening.

Our patch-based inference strategy—based on fixed-size regions (25.6 μm per side) without explicit single-cell segmentation—demonstrated robust performance even in wide-field images with cell confluency as high as 70% (Fig. 4a). Unlike conventional segmentation-dependent pipelines, which often fail under densely packed conditions due to overlapping cell boundaries and ambiguous contours^34^, our approach avoids reliance on accurate cell delineation and instead leverages local morphological cues for classification. This segmentation-free design enables scalable, high-throughput analysis in realistic imaging settings.

Excluding FasL + CHX + z-VAD–treated cells due to their ambiguous morphologies (Fig. 5c), the overall classification accuracy for A549 cells dropped to 58%. While apoptotic (81%) and drug-treated live cells (85%) were classified with reasonable accuracy, H_2_O_2_-induced necrotic cells were entirely misclassified (0%), and live-control cells were frequently confused with live-treated cells (only 29% correctly classified). These performance degradations likely stem from two key sources of domain mismatch. First, the model was trained exclusively on NaOH-induced necrosis in HeLa cells, which is characterized by rapid and homogeneous RI reduction. In contrast, oxidative necrosis in A549 cells triggered by H_2_O_2_ leads to more subtle, heterogeneous RI changes with granular cytoplasmic textures (Fig. 6). Second, the morphological distinction between control and treated A549 cells was less pronounced than in HeLa cells, and such fine-grained intra-class variability was not captured during model training.

Moreover, A549 cells treated with FasL + CHX + z-VAD displayed dynamic morphological transitions that deviated from canonical necroptosis. These cells often exhibited early apoptotic features—such as membrane blebbing and nuclear condensation—followed by late-stage membrane rupture and cytoplasmic dispersion. The lack of preserved nuclear architecture, a hallmark of necroptosis in HeLa cells, raises the possibility of secondary necrosis or atypical necroptotic programs. Without biochemical validation (e.g., MLKL phosphorylation or RIPK3 activation), these hybrid phenotypes could not be confidently classified based on morphology alone (Fig. 5a). This underscores the difficulty of phenotype interpretation under domain shift, where structural cues diverge from those in the training distribution.

## 4. Conclusion

We developed a deep learning framework integrated with 3D holotomography to enable real-time, label-free classification of major cell death pathways—apoptosis, necroptosis, and necrosis. By leveraging volumetric R) maps, our approach bypasses the need for fluorescence labeling or cell segmentation, allowing robust and scalable inference even in densely packed environments. The patch-wise strategy demonstrated high classification accuracy (97.2 ± 2.8%) in HeLa cells and maintained performance under high-confluency conditions where conventional segmentation-based methods often fail.

A key biological strength of our approach lies in its temporal sensitivity: the model successfully predicted necroptotic transitions several hours before conventional biochemical markers (Annexin V/PI) appeared, highlighting the potential of holotomography to detect early morphological signatures of regulated cell death.

We further evaluated generalizability by testing the model on A549 lung cancer cells—a domain-shifted context with distinct morphology and treatment response. The model exhibited substantial performance degradation, particularly in the classification of oxidative necrosis and live-control states. Cells treated with FasL + CHX + z-VAD displayed ambiguous, hybrid morphologies that were not adequately captured by the training distribution. These findings reveal that morphological cues alone may be insufficient to resolve complex or non-canonical phenotypes across diverse biological domains.

Overall, this study establishes holotomography-based AI as a technically powerful and biologically informative platform for high-resolution, label-free phenotyping of cell death. The segmentation-free design, early detection capability, and well-controlled evaluation across conditions position this framework as a valuable tool for high-throughput drug screening, cytotoxicity assessment, and mechanistic studies of cell fate. Future work should focus on incorporating biochemical markers, expanding training diversity, and applying domain adaptation techniques to further enhance generalizability and clinical relevance.

## 5. Methods

### 5.1 Cell culture and Maintenance

HeLa (ATCC CCL-2) and A549 (ATCC CCL-185) cells were cultured in Dulbecco’s Modified Eagle Medium (DMEM; ATCC 30-2002) supplemented with 10% fetal bovine serum and 1% penicillin– streptomycin (Thermo Fisher Scientific). Cells were maintained at 37°C in a humidified incubator with 5% CO_2_. For experiments, cells were seeded into TomoDishes (Tomocube Inc.) at a density of 1 × 10^5^ cells/mL and grown to approximately 70% confluency prior to treatment.

### 5.2 Cell Death Induction

Cell death was induced using recombinant human Fas ligand (100–200 ng/mL; Sino Biological, 10244-H07Y) crosslinked with anti-His antibody, or recombinant TNF-α (10–50 ng/mL; NKMAX, TNF0501) in combination with cycloheximide (CHX; 5–10 μg/mL). Sodium hydroxide (NaOH) and hydrogen peroxide (H_2_O_2_) were used to induce necrosis. Where indicated, cells were pre-treated with the pan-caspase inhibitor z-VAD-FMK (20–50 μM; Sigma, 627610) for 1 h to block apoptosis. HeLa cells were seeded and cultured for ∼18 h prior to treatment. All inducers were freshly prepared and applied at optimized concentrations. Imaging was conducted under sub-confluent conditions to minimize artifacts and allow accurate assessment of death phenotypes.

### 5.3 Fluorescence Staining

Cells were washed once with phosphate-buffered saline (PBS) and incubated with Hoechst 33342 (5 μg/mL; Thermo Fisher Scientific, H3570) at 37 °C for 15 min to label nuclei. Apoptotic cells were stained using the Alexa Fluor 488 Annexin V/Dead Cell Apoptosis Kit (Thermo Fisher Scientific, V35112) according to the manufacturer’s protocol. Fluorescence imaging was performed immediately using DAPI and FITC filter sets for Hoechst and Annexin V, respectively.

### 5.4 Imaging

Real-time holotomography imaging of cell death was performed using the HT-X1 system (Tomocube Inc.), which captures high-resolution time-lapse datasets at 30-minute intervals over a 24-hour period. The system employs four optimized illumination wavefronts to acquire transmitted intensity images, which are computationally reconstructed into three-dimensional RI tomograms via deconvolution using theoretically estimated point spread functions. Illumination was provided by a 449 nm center-wavelength LED, modulated by a DMD (DLP4500, Texas Instruments). The imaging optics included a condenser lens with a numerical aperture (NA) of 0.75 and an objective lens with an NA of 0.95.

### 5.5 Data preprocessing

The acquired imaging data were analyzed using a programming language python. Training of an AI model was done with the pytorch^35^ module, an open-source machine learning library in python.

Collected 3D holotomography images used in the data preprocessing were 16 hours after the start of imaging. The reason we chose 16 hours is because morphological features of each cell death pathways were stabilized at that point, so that time point are considered as representative cases for all pathways. After we collected the 3D images, we converted them into 2D MIP images. The 2D images were then divided into smaller patches. For example, if the 2D MIP image is 960 × 960 pixels and the patch size is 320 × 320 (approximately 51.2 μm per side), a total of nine patches are extracted from each image. Note that the patches don’t necessarily contain a whole single cell. It could contain part of the single cell, or multiple cells in one patch. Among the 2D patches, regions where cells don’t exist were excluded. The acquired 3D fluorescence images were also converted to 2D MIP for analysis.

### 5.6 Model Training

While we trained the model, we noticed that inclusion of the live (treated) class enhanced model classification accuracy. It could mean that the model can capture morphological differences when a cell is treated by drugs. Also, we adopted a fine-tuning strategy^36^ with a pre-trained model: ResNet-101. It is a model originally trained to classify 1000 different objects. By modifying the last few layers and training for the cell death data, the model succeeded at classifying cell death pathways with accuracy 97.2%.

Data augmentation is applied in the training step, such as cutmix^37^, flip, crop, resize and random noise. We used Adam optimizer^38^ and cross entropy loss function adjusted to the cutmix augmentation.

### 5.7 Model Inference Visualization

For inference, 3D holotomography images were first converted into 2D MIP images to retain key morphological features while reducing computational complexity. The resulting images were subdivided into fixed-size patches (160 × 160 pixels), which served as input units for classification.

To avoid misclassification of background regions—especially in sparse fields—we implemented a two-step patch filtering strategy. First, we identified patches where more than 95% of pixels exhibited refractive index (RI) values below 1.342, a threshold indicative of background. However, since necrotic cells also show decreased RI due to cytoplasmic leakage, this criterion alone was insufficient. To address this, we introduced a secondary classification using a lightweight convolutional neural network (ResNet-50) trained to distinguish necrotic cell patches from background and other death modalities. Patches predicted as background were excluded from downstream inference, while those classified as necrosis were retained.

For whole-field analysis, we employed a patch-overlapping inference strategy to ensure robustness and spatial continuity. The model slid over the full field-of-view (FOV) image, generating predictions for overlapping patches. Final class labels for each region were determined by majority voting among overlapping predictions, reducing noise and improving accuracy, particularly in high-confluency or heterogeneous regions.

## Data availability

All data are available from the corresponding authors upon reasonable request.

## Code availability

The code supporting this study is available via GitHub at https://github.com/papybara/cell_death Reference

## Acknowledgements

This work was supported by National Research Foundation of Korea grant funded by the Korea government (MSIT) (RS-2024-00442348, 2022M3H4A1A02074314), Korea Institute for Advancement of Technology (KIAT) through the International Cooperative R&D program (P0028463), and the Korean Fund for Regenerative Medicine (KFRM) grant funded by the Korea government (the Ministry of Science and ICT and the Ministry of Health & Welfare) (21A0101L1-12), Basic Science Research Program through the National Research Foundation of Korea (NRF) funded by the Ministry of Education (RS-2023-00241278). We thank Mahn Jae Lee, Chungnam National University, Korea, for helpful discussions on this study.

## Author Information

These authors contributed equally: Minwook Kim, Wei Sun Park.

## Authors and Affiliations

Department of Physics, Korea Advanced Institute of Science and Technology (KAIST), Daejeon, Republic of Korea

Minwook Kim, Wei Sun Park, Geon Kim, Juyeon Park & YongKeun Park

KAIST Institute for Health Science and Technology, Daejeon, Republic of Korea

Minwook Kim, Wei Sun Park, Geon Kim, Juyeon Park & YongKeun Park

Tomocube Inc., Daejeon, Republic of Korea

Sanggeun Oh, Jaephil Do, YongKeun Park

## Contributions

M.K. analyzed the data and developed the AI model. W.P., S.O., and J.D. designed and performed the experiments. M.K., W.P., and G.K. interpreted the results and contributed to data analysis. M.K. wrote the initial draft of the manuscript. All authors contributed to manuscript revision. Y.K.P. supervised the overall project.

